# Circular RNA circPLOD2 regulates pericyte function by targeting the transcription factor KLF4

**DOI:** 10.1101/2022.12.04.519017

**Authors:** Simone Franziska Glaser, Andre Brezski, Nina Baumgarten, Marius Klangwart, Andreas W. Heumüller, Ranjan Kumar Maji, Matthias S. Leisegang, Stefan Guenther, Christoph M. Zehendner, David John, Marcel H. Schulz, Kathi Zarnack, Stefanie Dimmeler

## Abstract

Circular RNAs (circRNAs) are generated by back-splicing and control cellular signaling and phenotypes. Pericytes stabilize the capillary structure and play an important role in the formation and maintenance of new blood vessels. Here, we characterized hypoxia-regulated circRNAs in human pericytes and showed that circPLOD2 is induced by hypoxia and regulates pericyte function. Silencing of circPLOD2 increased pericyte proliferation, endothelial-pericyte interaction and tube formation. Transcriptional profiling of circPLOD2-depleted cells and epigenomic analyses revealed widespread changes in gene expression and identified the circPLOD2-dependent regulation of the transcription factor KLF4 as a key effector of these changes. Importantly, overexpression of *KLF4* was sufficient to reverse the effects on pericyte proliferation and endothelial-pericyte interactions observed after circPLOD2 depletion. Together, these data revealed a novel function of circPLOD2 in the control of pericyte proliferation and capillary formation and showed that circPLOD2-mediated regulation of KLF4 significantly contributes to the transcriptional response to hypoxia.

**Highlights:**

- circPLOD2 is upregulated in hypoxia in human vascular pericytes
- Selective depletion of circPLOD2, but not linear *PLOD2* mRNA, changes pericyte migration and endothelial-pericyte interaction
- circPLOD2 depletion triggers widespread changes in gene expression that are mirrored in the transcriptional hypoxia response
- Epigenomic analyses pinpoint the transcription factor KLF4 as a central player in circPLOD2-mediated expression changes
- *KLF4* overexpression is sufficient to rescue the changes in pericyte function caused by circPLOD2 depletion

## Introduction

The vasculature plays an important role in maintaining oxygen and nutrient supply. Tissues with high oxygen demand, such as the heart or brain, are highly vascularized by small vessels termed capillaries. These are formed by endothelial cells, which are surrounded by pericytes. Endothelial cells and pericytes are connected to each other by gap junctions and further interact by paracrine and multiple ligand-receptor complexes^1^. Pericytes stabilize the capillary structure and support the formation of new blood vessels. Injury or dysfunction of pericytes lead to destabilization of the capillaries and can results in pathologies, such as diabetic retinopathy. Pericytes additionally fine-tune blood supply for example in the brain, since they can control the diameter of the capillaries by inducing contractions^2^. During ischemic stroke, pericytes intervene at multiple stages and contribute to the acute stress response as well as to adaptation and recovery^3^.

Circular RNAs (circRNAs) are a specialized form of RNAs that are broadly expressed in many tissues and can perform a variety of functions^4–6^. They are generated via a process termed back-splicing, in which the 3′ splice site of an exon is joined to a downstream 5′ splice site of the same or downstream exon(s), thereby forming a covalently closed circle. circRNAs usually originate from protein-coding genes that also produce linear transcript isoforms. Most circRNAs are exported to the cytoplasm^4^, where some can be translated, while most are thought to act as noncoding RNAs (reviewed in ^7^). Several mechanisms of action have been postulated both in the nucleus and the cytoplasm. Among these, microRNA (miRNA) sponging was initially described as a major role, however, this function appears to be restricted to a few circRNAs with multiple miRNA binding sites such as CDR1as (also known as CiRS-7)^6,8^. circRNAs also can act as scaffolds for RNA-binding proteins or affect transcription and host gene expression directly^9^.

The expression of circRNAs is highly dynamic and regulated between tissues and physiological conditions. They are particular prominent in the brain where they appear as an integral part of the transcriptome^10^. Moreover, they are strongly induced by various stimuli, including hypoxia, to promote the cellular response and adaption^11,12^. For instance, it was shown that circZNF292 is induced in endothelial cells under hypoxia and controls endothelial cell functions by interacting with the RNA-binding protein SDOS to strengthen cytoskeletal protein interaction^13,14^. circRNAs thus play important roles in the control of the vasculature. However, most work has been done in endothelial cells, while only few studies assessed their role in pericytes, mainly in the context of diabetes. In a mouse model for diabetic-induced stress conditions, it was found that the circRNA circPWWP2A is important for the pericyte-endothelial cell crosstalk and prevents diabetes-induced microvascular dysfunction^15^, and that circZNF532 protects against diabetes-induced retinal pericyte degeneration and vascular dysfunction^15^. However, the role of circRNAs in other pericyte functions, particularly in the context of ischemia, remains elusive.

Here, we characterize the expression and regulation of circRNAs in human vascular pericytes under hypoxia to mimic the declined oxygen supply by myocardial infarction or stroke. We identify multiple circRNAs that are induced in response to hypoxia, most prominently circPLOD2 which we decided to study further. While this circRNA had been previously shown to enhance the proliferation of ovarian cancer cells^15^, its role in hypoxia remains unknown. Intriguingly, we overserve that selective depletion of circPLOD2, but not linear *PLOD2* transcripts, improves endothelial-pericyte crosstalk and tube formation. Transcriptional profiling and epigenomic analyses reveal that circPLOD2 depletion triggers widespread changes in gene expression that result from regulation of the transcription factor KLF4 and are mirrored in the hypoxia response when circPLOD2 is strongly upregulated. Importantly, we demonstrate that KLF4 overexpression is sufficient to rescue the changes in pericyte function caused by circPLOD2 depletion.

## Results

### circPLOD2 is induced upon hypoxia in human pericytes

To determine the expression and regulation of circRNAs in pericytes, we used RNA sequencing (RNA-seq) data of rRNA-depleted total RNA from human brain vascular pericytes^16^. The cells had been exposed to 1% O_2_ and 5% CO_2_ for 24 hours (h) to mimic ischemia conditions, as they can occur in brain or hearts after infarction. Differential gene expression analysis showed an induction of 1,385 genes, which include the typical hypoxia-induced gene *vascular endothelial growth factor A* (*VEGFA)* (**Figure 1A**), whereas 1,393 genes were repressed (**Figure S1A, Table S1**). Differentially expressed genes (DEGs) were associated with gene ontology (GO) terms related to the response to oxygen levels and metabolic processes (**Figure 1B**), confirming typical hypoxia responses in pericytes.

**Figure 1.**
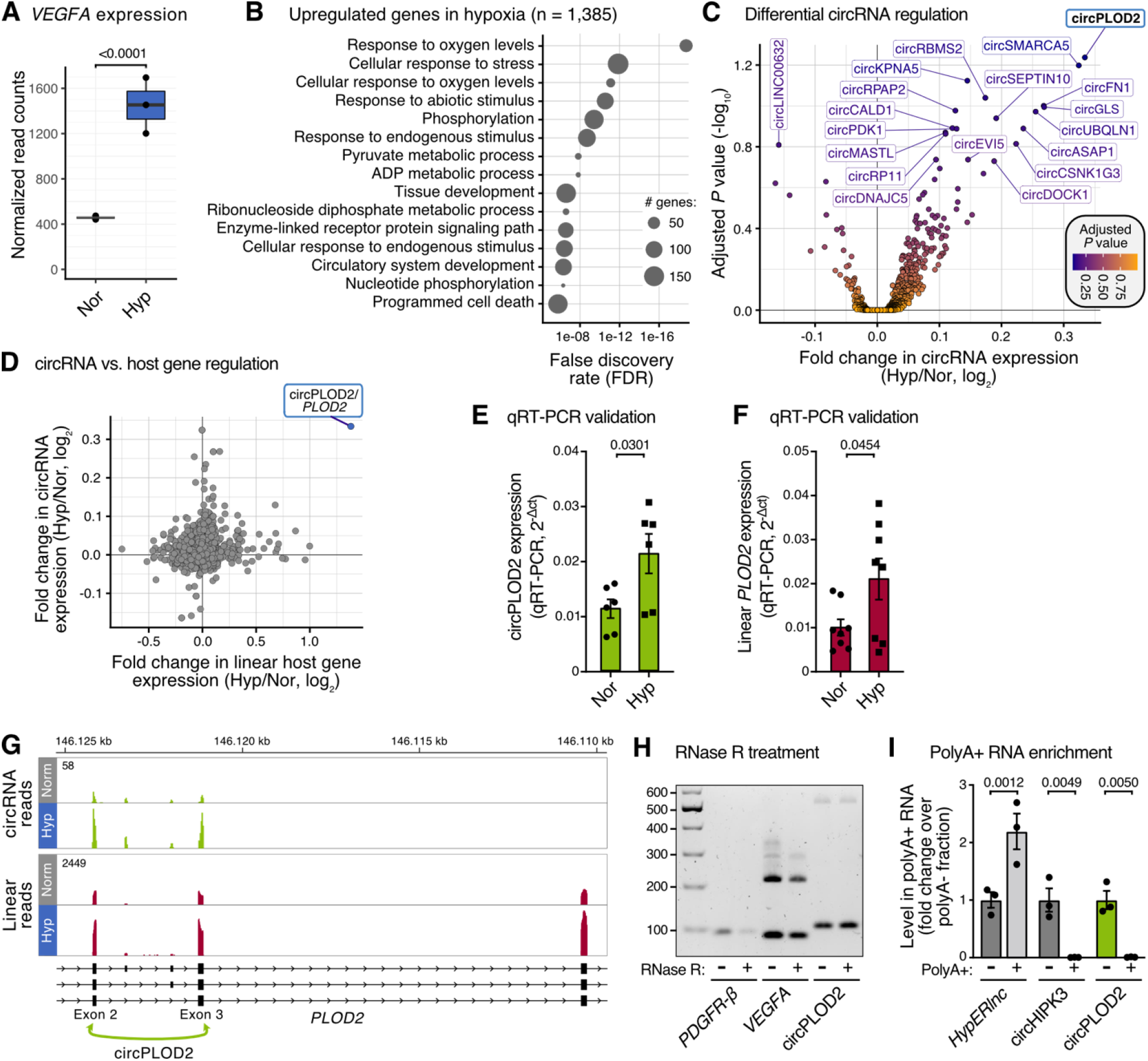
circPLOD2 is upregulated in human pericytes under hypoxia. **(A)** *VEGFA* mRNA is induced in pericytes under hypoxia. Human brain vascular pericytes were grown in normoxia (Nor; 21% O_2_, 5% CO_2_) or hypoxia (Hyp; 1% O_2_, 5% CO_2_) for 24 h. **(B)** Pericytes show a typical transcriptional hypoxia response. Top 15 gene ontology (GO) Biological Process terms enriched in hypoxia-upregulated genes (n=1,385). **(C)** Multiple circRNAs are upregulated in hypoxic pericytes. Volcano plot shows log_2_-transformed fold change in circRNA expression (Hyp/Nor) against adjusted *P* value (Benjamini-Hochberg correction). circRNAs with adjusted *P* value<0.2 are labeled. **(D)** circPLOD2 increase is mediated by concomitant upregulation of the linear *PLOD2* gene. Scatter plot compares log_2_-transformed fold change in circRNA and linear host gene expression (Hyp/Nor) for 946 circRNAs with sufficient read counts. **(E-F)** Validation of circPLOD2 (E) and linear PLOD2 (F) regulation upon hypoxia (Hyp) or normoxia (Nor) on mRNA level, measured by qRT-PCR. Data were normalized to *RPLP0* and visualized as relative gene expression (2-ΔCt; n=6-8). **(G)** circPLOD2 is generated via back-splicing of exon 2 and 3 of *PLOD2*. Genome browser view shows RNA-seq coverage for circPLOD2 (back-splice reads, green) and linear *PLOD2* (red) on exons 2-4 of the *PLOD2* gene (complete gene shown in **Figure S2A**). **(H)** circPLOD2 is RNase R-resistant. Representative gel showing PCR products on total RNA from pericytes with RNase R digestion (RNase R +) or without (RNase R -). circPLOD2 was amplified with specific primers across the back-splice junction. Linear *PDGFR-β* and *VEGFA* mRNAs were used as controls (n=2). **(I)** circPLOD2 is present in poly(A)− RNA fraction. qRT-PCR quantification of circRNAs circPLOD2 and circHIPK3 after poly(A)+/− RNA fractionation using oligo-dT beads. lncRNA *HypERlnc* served as polyadenylated positive control. Data are depicted as fold change over poly(A)− fraction (n=3). Data are shown as mean ± standard error of mean (s.e.m.) **(E, F, I)**. Normal distribution was assessed using the Shapiro-Wilk test. Statistical analysis to compare two groups was performed using unpaired, two-sided Student’s *t*-tests **(E, F)** or more groups with 2-way ANOVA with multiple comparison and Bonferroni correction **(I)**.

To analyze the circRNA response in pericytes, we quantified circRNAs across all samples using our computational pipeline Calcifer (see Methods, **Figure S1B**). In total, we identified 6,187 circRNAs which primarily originated from protein-coding genes (**Figure S1C, Table S2**). Despite the typically low RNA-seq read counts for circRNAs (as only reads spanning the back-splice junction can be unambiguously assigned to the circular isoform of a transcript), we were able to detect several circRNAs that apparently responded to the hypoxia treatment in pericytes (**Figure 1C, Figure S1D, Table S2**). The vast majority of the hypoxia-responsive circRNAs were upregulated in hypoxia. Among these, the most significantly and strongly upregulated circRNA was circPLOD2 (**Figure 1C**). The upregulation of circPLOD2 in hypoxic pericytes was achieved by a concomitant upregulation of the *PLOD2* host gene (**Figure 1D**). We confirmed the significant induction of circPLOD2 and the underlying linear *PLOD2* transcript in hypoxic pericytes using quantitative real time (qRT)-PCR (**Figure 1E, F**).

The circRNA circPLOD2 is generated via back-splicing from exon 3 to exon 2 of the *PLOD2* pre-mRNA (**Figure 1G, Figure S2A**). While the major isoform that was primarily induced upon hypoxia included only exons 2 and 3, we also detected two additional isoforms that resulted from the occasional inclusion of two intervening, noncoding exons between the back-splice sites (**Figure 1G, Figure S2A, B**). The exact positioning of the back-splice junction between exons 3 and 2 was confirmed by Sanger sequencing (**Figure S2C**). We further demonstrated that circPLOD2 was resistant to the 3′-5′ exonuclease RNase R (**Figure 1H**) and was not detected in poly(A)-selected RNAs (**Figure 1I**), confirming the circular nature of this RNA. We concluded that circPLOD2 is a circular isoform of *PLOD2* that is strongly upregulated in hypoxic pericytes.

### Depletion of circPLOD2 augments endothelial-pericyte interactions and vascular tube formation

To investigate the function of circPLOD2 in human pericytes, we selectively depleted circPLOD2 using siRNAs directed against the back-splice junction between *PLOD2* exons 3 and 2. qRT-PCR confirmed that the siRNA treatment efficiently reduced the circular RNA circPLOD2 but did not affect the expression of the linear host transcript (**Figure 2A, B**). Of note, we found that migration of circPLOD2-depleted pericytes, measured in the Boyden chamber assay, was significantly increased (**Figure 2C**). In addition, depletion of circPLOD2 significantly promoted the incorporation of bromodeoxyuridine (BrdU), indicative of active DNA replication (**Figure 2D**), suggesting that circPLOD2 influences the migration and proliferation of pericytes.

**Figure 2.**
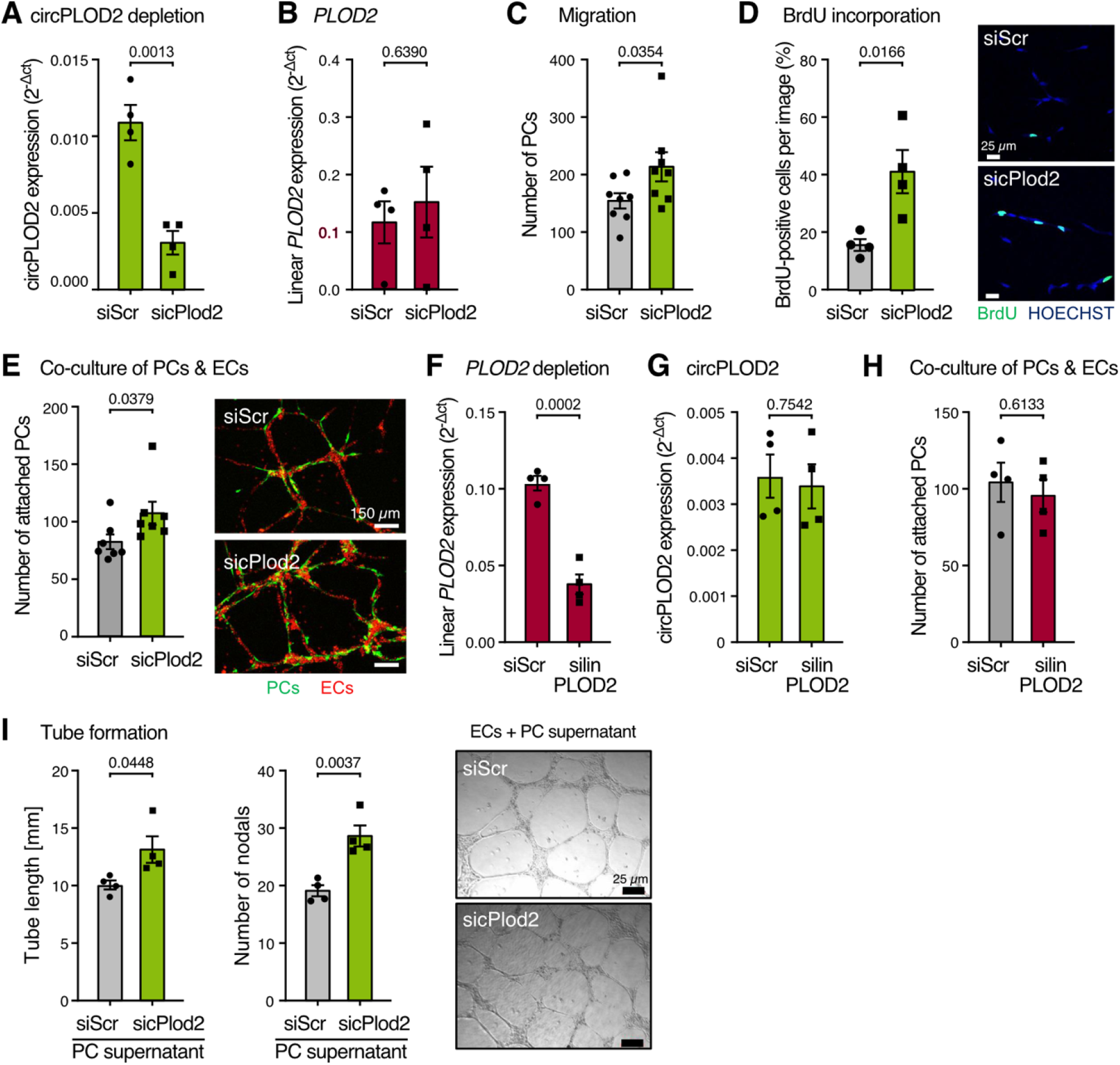
circPLOD2 depletion impairs pericyte function. Non-targeting siRNA (siScr) or siRNAs targeting circPLOD2 (sicPLOD2), linear *PLOD2* (silinPLOD2) were transfected into pericytes and the function was assessed. **(A, B)** circPLOD2 is efficiently depleted using siRNAs. qRT-PCR quantifications of circPLOD2 (A) and linear *PLOD2* transcripts (B) in pericytes. Data were normalized to *RPLP0* and visualized as relative gene expression (2^-ΔCt^; n=4). **(C)** Migration of pericytes through a Boyden chamber was determined by counting DAPI-stained cells after siRNA-mediated circPLOD2 depletion with ImageJ (n=8, 5 images per condition). **(D)** Proliferation of pericytes was assessed after 6 h incubation with bromodeoxyuridine (BrdU) by counting BrdU-positive cells, normalized to all cells per image. Quantification is shown in the left panel, with representative images on the right (n=4, 3 images per condition). Nuclei were stained with HOECHST, depicted in blue and BrdU is shown in green (scale: 25 µm). **(E)** Pericyte-endothelial cell co-culture assay after circPLOD2 depletion. Pericytes were labelled with GFP (green), and endothelial cells stained with Dil-LDL (red). Pericytes which were attached to the endothelium were counted per image. Quantification is shown in the left panel, with representative images on the right (n=7, 3-8 images per condition; scale bar: 150 µm). **(F, G)** siRNA-mediated depletion of linear *PLOD2*, but not circPLOD2, in pericytes was validated on RNA level, by measuring linear *PLOD2* (F) and circPLOD2 (G) by qRT-PCR. Data were normalized to *RPLP0* and visualized as relative gene expression (2^-ΔCt^; n=4). **(H)** Pericyte-endothelial cell co-culture assay after depletion of linear *PLOD2*. Labelling as in (E). Pericytes which were attached to the endothelium were counted per image (n=4, 3-6 images per condition; scale bar: 150 µm). **(I)** Tube formation assay of endothelial cells incubated with supernatant of siRNA-treated pericytes (siScr or sicPlod2). Quantification of endothelial tube length in mm is shown on the left side, left panel, and number of nodals per image is shown in the right panel. Representative images are shown on the right side (n=4, 3-4 images per condition; scale bar: 25 µm). All data are shown as mean ± s.e.m.. Normal distribution was assessed using Shapiro-Wilk test. Statistical analysis to compare two groups was performed using unpaired, two-sided Student’s *t*-tests **(A, B, D, F-H)** or Mann-Whitney test **(C, E)**. Outliers were identified with the Grubbs Outlier test **(C)**.

To test for effects on pericyte function, we co-cultured tracked circPLOD2-depleted and control pericytes with endothelial cells and monitored the interaction of both cell types to form vessel-like structures (**Figure 2E**). Importantly, depletion of circPLOD2 in pericytes significantly increased the number of pericytes which attached to the unmodified endothelial cells (**Figure 2E**), indicating a role in endothelial-pericyte crosstalk. To address whether these effects are unique to the circRNA, we designed specific siRNAs to silence the linear *PLOD2* transcript, but not circPLOD2, by targeting exon 17 of *PLOD2* (**Figure 2F, G**). Targeted depletion of the linear *PLOD2* transcript did not affect pericyte attachment to endothelial cells (**Figure 2H**), supporting that circPLOD2 has a circRNA-specific function in endothelial-pericyte interaction. To address whether circPLOD2 directly interferes with paracrine signaling from pericytes to endothelial cells, we tested the pro-angiogenic capacity of supernatant obtained from circPLOD2-depleted and control pericytes. Indeed, supernatant of circPLOD2-depleted pericytes was sufficient to augment tube formation in monocultures of endothelial cells (**Figure 2I**), suggesting that circPLOD2 regulates soluble factors which promote endothelial cell tube formation. Altogether, we could demonstrate that circPLOD2 depletion impairs pericyte function at several levels, including cell motility, endothelial-pericyte crosstalk via paracrine signaling and the stimulation of endothelial tube formation.

### circPLOD2 modulation causes widespread changes in gene expression which also occur in hypoxia

To understand which changes occur in pericytes upon circPLOD2 depletion, we performed RNA-seq of circPLOD2-depleted and control pericytes (n = 3 biological replicates, **Figure 3A**). Calcifer quantification confirmed the depletion of circPLOD2 (**Figure 3B**). Strikingly, circPLOD2 depletion caused massive changes in gene expression, as 2,510 genes significantly changed in expression, including 1,450 upregulated and 1,060 downregulated genes (**Figure 3C, Table S3**). GO analysis revealed that the upregulated genes included many genes associated with cell cycle progression and proliferation (**Figure 3D, E**), which was consistent with the increase in BrdU incorporation in circPLOD2-depleted pericytes (**Figure 2D**).

**Figure 3.**
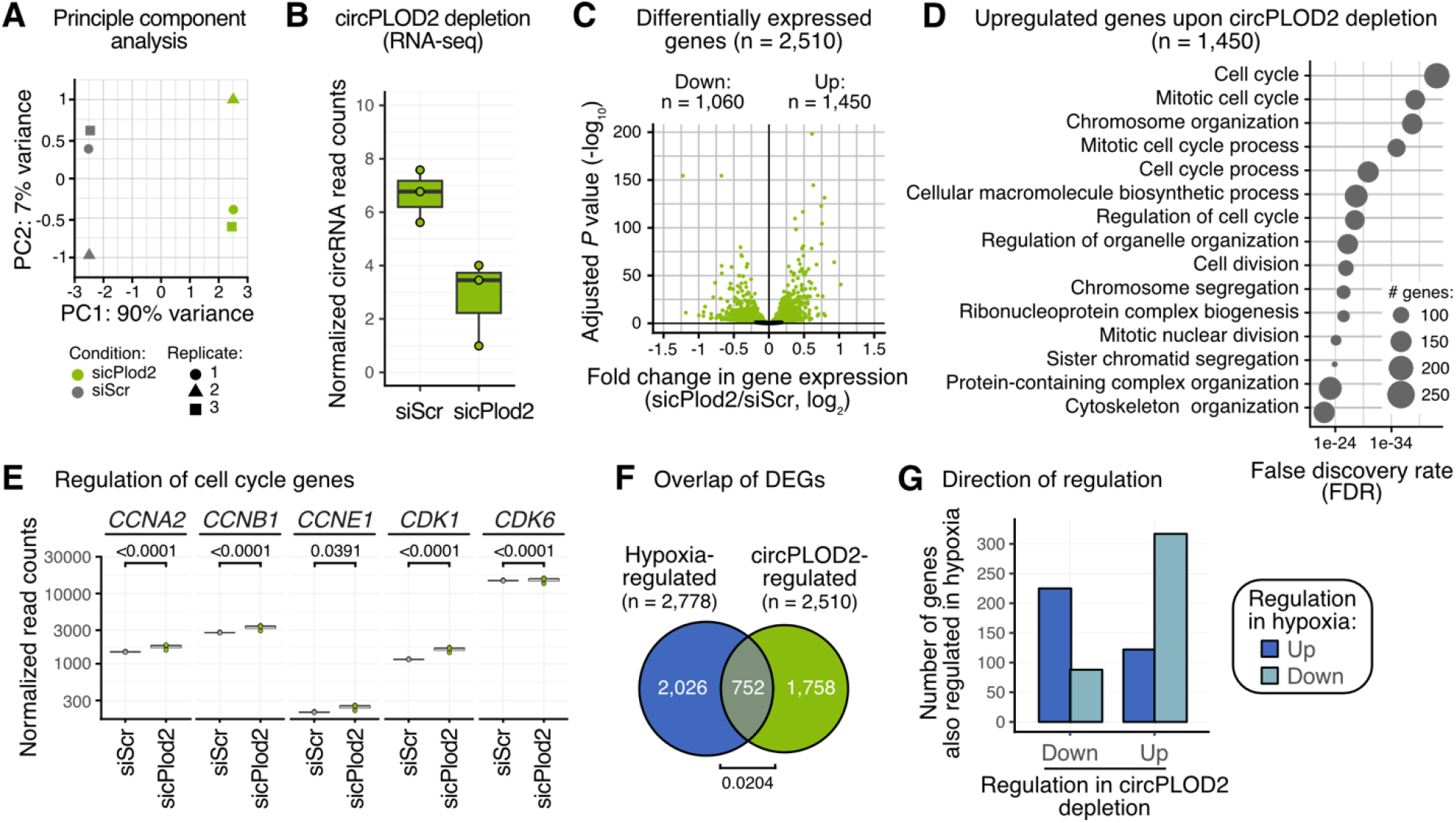
circPLOD2 depletion causes the regulation of hundreds of genes which inversely respond to the circPLOD2 upregulation in hypoxia. **(A)** Principal component analysis (PCA) shows distinction of circPLOD2-depleted (sicPlod2) and control (scrambled siRNA, siScr) pericytes (n=3). PCA was performed on 500 genes with highest variance using DESeq2. **(B)** circPLOD2 is efficiently depleted in the RNA-seq data. Boxplot shows compares number of back-splice reads (normalized to library size) in RNA-seq replicates. **(C)** circPLOD2 depletion causes widespread changes in gene expression. Volcano plot shows log_2_-transformed fold change in gene expression (Hyp/Nor) against adjusted *P* value. Significantly regulated genes are shown in green (adjusted *P* value<0.1, Benjamini-Hochberg correction). **(D)** Upregulated genes are associated with cell cycle progression and proliferation. Top 15 GO Biological Process terms enriched in genes upregulated upon circPLOD2 depletion (n=1,450). **(E)** RNA-seq read counts (normalized to library size) for selected cell cycle-associated genes. **(F)** Hypoxia-regulated genes are significantly enriched for circPLOD2-regulated genes. Venn diagram shows overlap between differentially expression genes (DEGs) in response to hypoxia (blue) or circPLOD2 depletion (green, *P* value from Fisher’s exact test). **(G)** Differentially expressed genes are inversely regulated upon circPLOD2 depletion and hypoxia where circPLOD2 is induced. Barchart indicates number of genes that are up- or downregulated in hypoxia and circPLOD2 depletion.

To test whether the circPLOD2-regulated genes also respond in hypoxia, we overlaid our RNA-seq results from both experiments. Indeed, we observed a significant overlap of differentially expressed genes changing upon hypoxia or circPLOD2 depletion in pericytes (**Figure 3F**). Consistent with the upregulation of circPLOD2 in hypoxia, the majority of genes showed an inverse effect compared to the circPLOD2 depletion, i.e., genes that went down upon circPLOD2 depletion were preferentially upregulated in hypoxia, and vice versa (**Figure 3G, S3A**). Together, these observations suggest that circPLOD2 induction in hypoxia causes widespread changes in gene expression which substantially contribute to the overall hypoxia response in pericytes.

### Changes in gene expression are mediated via regulation of the transcription factor KLF4

Intrigued by the substantial contribution of circPLOD2-mediated regulation to the transcriptional hypoxia response of human pericytes, we investigated which pathway could mediate the gene expression changes observed upon circPLOD2 depletion.

We first looked at potential miRNA involvement. However, contrary to a previous report^17^, we observed neither a generally increased density of miRNA target sites nor an enrichment for any particular miRNA (**Figure S3B**). Moreover, in publicly available AGO2 pulldown data^18^, we only found two peaks of AGO2 binding in circPLOD2, presumably harboring two miRNAs, however, their predicted targets showed no directed regulation in circPLOD2-depleted pericytes (**Figure S3C**). Finally, target sites of the previously described circPLOD2-interacting miRNA miR-378^17^ were neither predicted nor detected in the AGO2 pulldowns. Based on these observations, we concluded that circPLOD2 most likely does not interfere with miRNA regulation.

Instead, the extensive changes in gene expression upon circPLOD2 depletion made us suspect that one or more transcription factors (TFs) may be deregulated. To test this, we integrated our expression data with epigenomic information to search for transcriptional regulators that could mediate the observed changes. To investigate which TFs might be affected, we collected regulatory elements (REMs) associated to the top 1,000 DEGs after circPLOD2 depletion. REMs, such as enhancers or repressors, are genomic regions which harbor TF binding sites and show cell type-specific variability in chromatin accessibility to regulate gene expression^19,20^. The REMs were downloaded from the EpiRegio database^20^, which is a comprehensive resource holding more than 2.4 million REMs linked to human target genes computed from paired DNase-I-seq and RNA-seq data from several cell types^21^. We used published DNase I sensitivity profiles collected from ENCODE^20^ (see Methods) to determine which REMs are active in human pericytes (mean DNase I signal > 0.1). The resulting 68,086 REMs were used to perform a TF motif enrichment analysis using the rank-based PASTAA algorithm^22^ and known TF binding motifs for 292 TFs that were expressed in our RNA-seq data (transcripts per million [TPM] > 1). We observed a significant enrichment of 95 TFs (*P* value≤0.01, Benjamini-Hochberg correction) in the REMs of the top 1,000 DEGs upon circPLOD2 depletion.

To identify the TFs which most likely have the largest effect on differential expression, we counted the number of predicted binding sites for each significantly enriched TF for all REMs (**Figure 4A**). Several members of the Krüppel like factor (KLF) family, among them KLF4, showed multiple TF binding sites in a large number of REMs. Among these, KLF4 stood out in particular as it had the second highest number of associated REMs (n = 3,025 active REMs with a KLF4 binding site associated with 767 DEGs out of the top 1,000 DEGs; **Figure 4B**) and it was significantly down-regulated after circPLOD2 depletion (**Figure 4C**). DNase I sensitivity profiles confirmed that the KLF4-associated REMs at DEGs showed an open chromatin status in pericytes under control conditions (**Figure 4D**), supporting that a KLF4-dependent regulation of these genes might occur. Consistent with a role of KLF4 in the transcriptional changes seen upon circPLOD2 depletion, as well as upon hypoxia where circPLOD2 is induced, expression of *KLF4* was inversely regulated in both treatments (**Figure 4E, F**). We further validated the downregulation of *KLF4* expression upon circPLOD2 depletion by qRT-PCR (**Figure 4G**).

**Figure 4.**
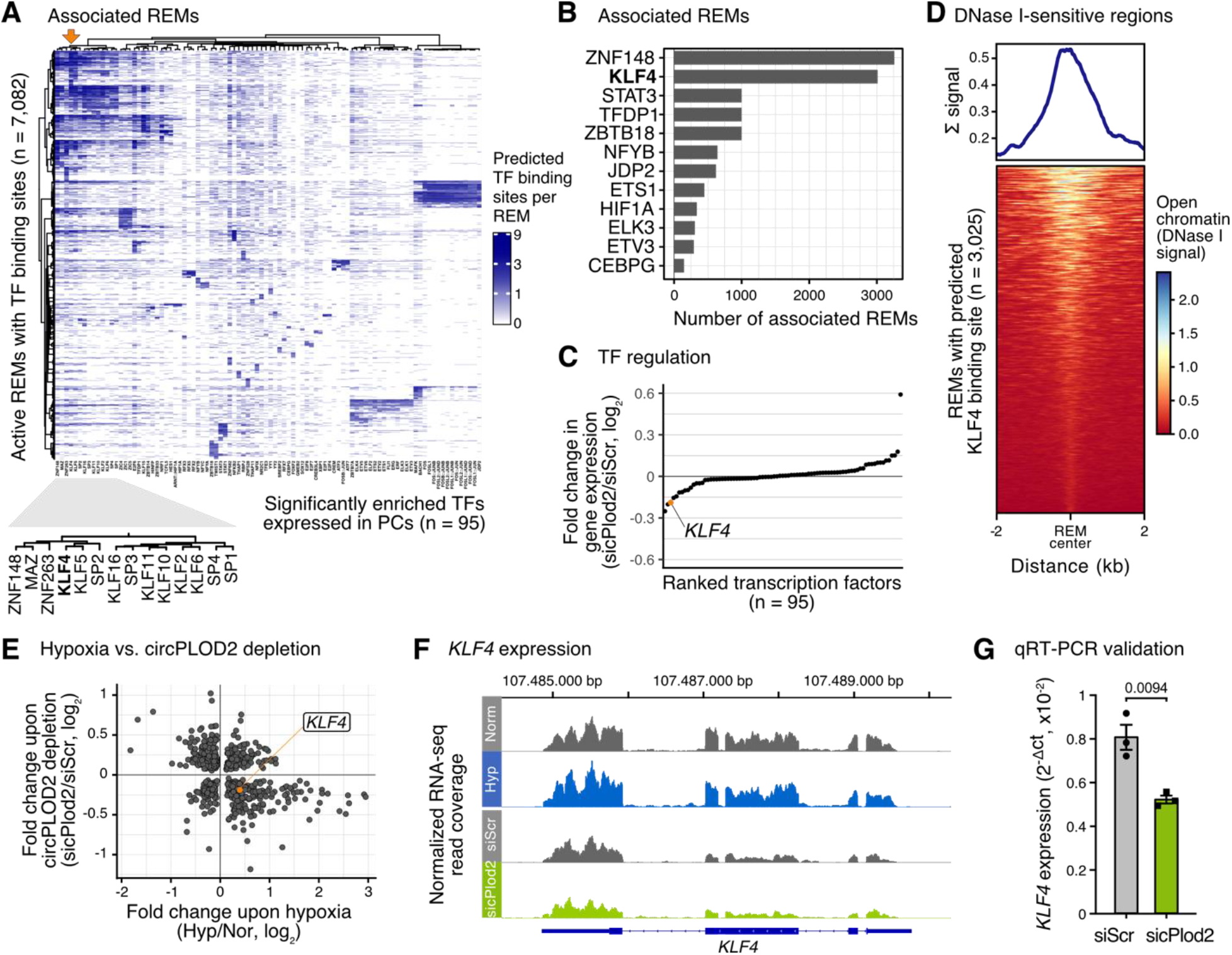
Gene expression changes result from KLF4 downregulation upon circPLOD2 depletion. **(A)** Heatmap shows number of predicted transcription factor (TF) binding sites per regulatory element (REM; y-axis) for the 95 significantly enriched TFs (x-axis). Color scale indicates number of TF binding sites per REM. Zoom-in shows cluster enriched for Krüppel like factor (KLF) family, including KLF4 (bold). **(B)** Barplot visualizes 12 TFs with the highest number of associated REMs, i.e., harboring least one predicted binding site of the TF. **(C)** Rank plot shows log_2_-transformed fold changes in gene expression upon circPLOD2 depletion (DESeq2) for mRNAs encoding 95 significantly enriched TFs. *KLF4* mRNA is highlighted in orange. **(D)** DNase I sensitivity signal for REMs with predicted KLF4 binding sites. Metaprofile (top) and heatmap (bottom) show a 4-kb window around REM centers. Color scale indicates chromatin accessibility. **(E, F)** *KLF4* mRNA is inversely regulated upon hypoxia (x-axis) and circPLOD2 depletion (y-axis). **(E)** Scatter plot compares shrunken, log_2_-transformed fold changes from DESeq2. (**F**) Genome browser view shows normalized RNA-seq coverage (counts per million) for *KLF4* in pericytes in normoxia (Nor) and hypoxia (Hyp, blue) as well as after circPLOD2 depletion (sicPlod2, green) vs. control (siScr). **(G)** *KLF4* mRNA expression in circPLOD2-depleted and control pericytes, measured qRT-PCR. Data were normalized to *RPLP0* and visualized as relative gene expression (2^-ΔCt^; n=3) and shown as mean ± s.e.m.. Normal distribution was assessed using the Shapiro-Wilk test. Statistical analysis to compare two groups was performed using unpaired, two-sided Student’s *t*-tests. Outliers were identified with the Grubbs Outlier test.

To gain further insights into how circPLOD2 may interfere with *KLF4* expression, we compared mRNA stability after blocking transcription with actinomycin D (ActD) in circPLOD2-depleted and control pericytes. As expected, *KLF4* mRNA was rapidly degraded within 60 minutes after ActD addition (**Figure 5A**). Of note, circPLOD2 depletion further reduced *KLF4* mRNA abundance (**Figure 5A**), indicating that the *KLF4* transcripts were significantly destabilized in absence of circPLOD2. Together, these data suggested that circPLOD2 stabilizes *KLF4* mRNA.

**Figure 5:**
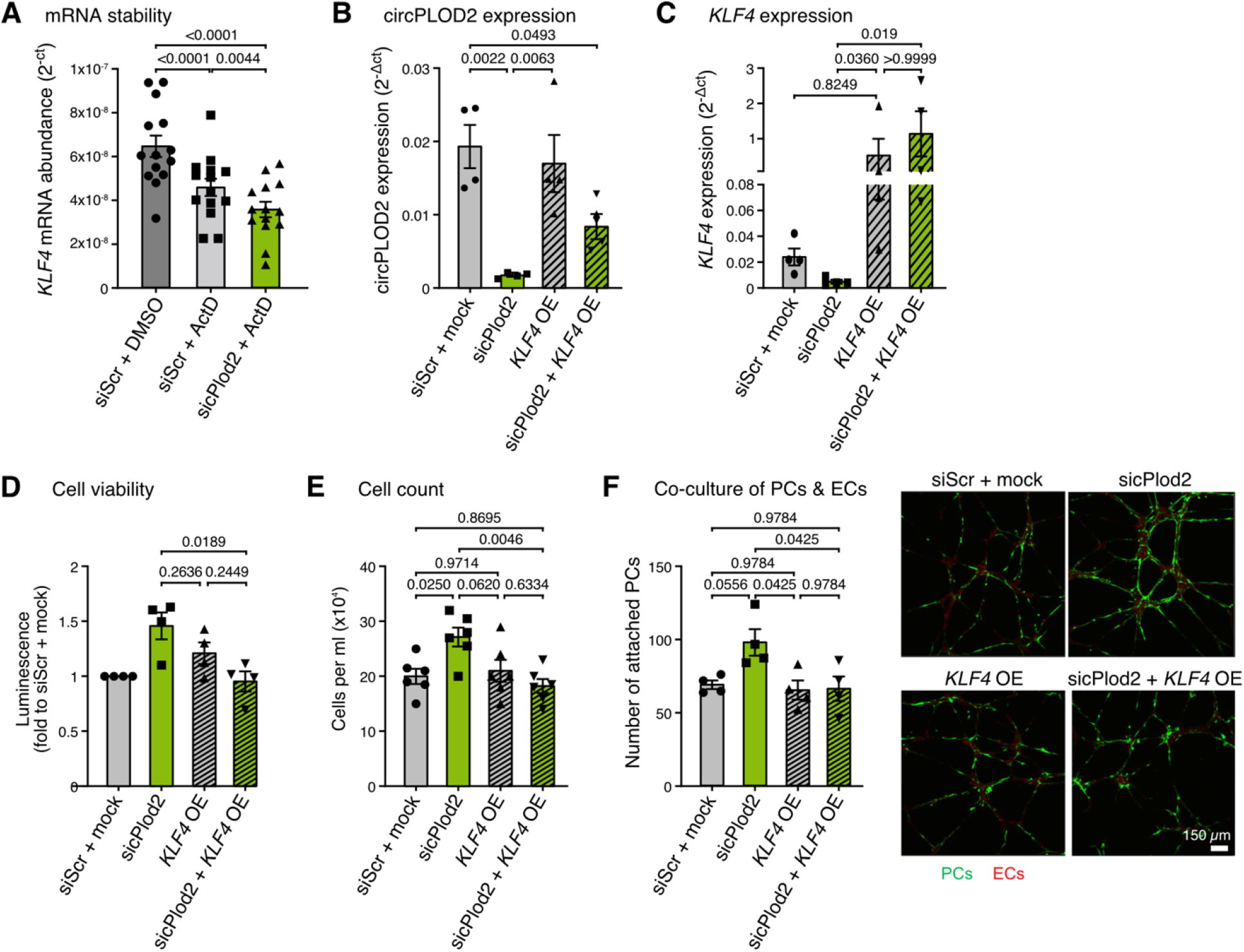
KLF4 overexpression is sufficient to restore pericyte function under circPLOD2 depletion. **(A)** *KLF4* mRNA abundance in circPLOD2-depleted and control pericytes treated with actinomycin D (ActD) or DMSO for 1 h. Data are shown as 2^-Ct^ values (n=14). **(B-F)** Pericytes were co-transfected with combinations of non-targeting siRNAs (siScr) and mock plasmids or siRNAs targeting circPLOD2 (sicPLOD2) and mock plasmids or *KLF4* overexpression (OE) with siScr. **(B, C)** Validation of circPLOD2 depletion and *KLF4* overexpression. Expression of circPLOD2 **(B)** and *KLF4* **(C)** was measured by qRT-PCR. Data were normalized to *RPLP0* and visualized as relative gene expression (2^-ΔCt^; n=4). **(D)** Cell viability of siRNA- and/or plasmid-transfected pericytes was determined by measuring the luminescence (relative luminescence units, RLU). Data are depicted as fold change over control (n=4). **(E)** Transfected pericytes were counted after trypsination. Cell number is depicted as cells per ml (*10^4^) based on a growth area of 21 cm^2^. **(F)** Pericyte-endothelial cell co-culture assay after circPLOD2 depletion and *KLF4* overexpression. Pericytes were labelled with GFP (green), and endothelial cells stained with Dil-LDL (red). Pericytes which were attached to the endothelium were counted per image. Quantification is shown in the left panel, with representative images on the right (n=4, 3-5 images per condition; scale bar: 150 µm). All data are shown as mean ± s.e.m.. Normal distribution was assessed using the Shapiro-Wilk test. Statistical analysis to compare two groups was performed using one-way ANOVA with multiple comparison and Tukey correction **(A, C, D, E)**, paired ANOVA with multiple comparison and Tukey correction **(F)**, Kruskal-Wallis test **(B)**. Outliers were identified with the Grubbs Outlier test **(E, F)**.

KLF4 has already been described to have important functions in pericyte and related smooth muscle cell biology^23,24^. To determine whether KLF4 downregulation upon circPLOD2 depletion might contribute to the observed biological effects, we used vectors to overexpress *KLF4* in circPLOD2-depleted pericytes. qRT-PCR confirmed that we achieved an almost 50-fold overexpression of *KLF4* in circPLOD2-depleted and control pericytes, and that *KLF4* overexpression did not interfere with circPLOD2 depletion (**Figure 5B, C**). Importantly, *KLF4* overexpression was sufficient to completely prevent the effects of circPLOD2 depletion: Cell viability, cell numbers and endothelial-pericyte attachment returned to control levels when both treatments were combined (**Figure 5D-F**). Together, these data demonstrate that restoring *KLF4* expression is sufficient to rescue circPLOD2 depletion in these assays. They further suggest that the circPLOD2-mediated downregulation of KLF4 significantly contributes to the physiological effects induced by circPLOD2 depletion.

## Discussion

circRNAs have been reported to play important roles in multiple tissues including the vasculature^15^. Given the limited insights into circRNA functions in human pericytes, we set out to characterize the circRNA repertoire and its response to hypoxia as a model for ischemic stress. In total, we detected more than 6,000 circRNAs in human vascular pericytes that mostly originated from protein-coding genes. In line with reports from other cell types^11,13^, we found that hypoxia induces the expression of multiple circRNAs. The most strongly upregulated circRNA was circPLOD2, which had already been described as hypoxia-regulated circRNA in cancer cells^11^. Upregulation of circPLOD2 was achieved by a concomitant increase in the host gene *PLOD2* which is a known target of the hypoxia-induced transcription factor HIF-1^25^. *PLOD2* encodes for a procollagen-lysine,2-oxoglutarate 5-dioxygenase that catalyzes the hydroxylation of lysyl residues in collagen-like peptides (reviewed in^26^) and thereby regulates the stability of intermolecular crosslinks within the extracellular matrix^25^.

Importantly, using a selective depletion of the circular isoform, we could demonstrate that circPLOD2 exhibits specific functions that were not seen for the linear host gene in pericytes. We found that circPLOD2 depletion augments cell proliferation, endothelial-pericyte interaction and pericyte migration, suggesting that its downregulation supports the formation of new capillaries. Inversely, the strong upregulation of circPLOD2 upon hypoxia may contribute to a limitation of endothelial-pericyte interactions under pathological conditions. Even though the linear transcript *PLOD2* is likewise upregulated upon hypoxia, the depletion of linear *PLOD2* had no effect on endothelial-pericyte interaction, highlighting a specific role for circPLOD2 in pericyte functionality. In endothelial cells, hypoxia is well known to induce many pro-angiogenic factors such as VEGF, but it also augments anti-angiogenic and cytotoxic proteins, which impair the recovery of the vasculature (reviewed in^22^). Indeed, previous studies support a detrimental role of hypoxia on vascular pericytes: for instance, preventing the hypoxia response in pericytes by HIF-1 loss-of-function preserved barrier integrity by reducing pericyte death and thereby maintaining vessel coverage and junctional protein organization^27^. Although we lack *in vivo* evidence, our *in vitro* findings are consistent with a model that the strong induction of circPLOD2 may contribute to the disturbed vessel stability under hypoxia by regulating pericyte functions.

Following the well-documented showcase example of the circRNA cDR1as, which regulates the levels of miR-7^6,8^, many studies reported purported functions of circRNAs in the sponging of miRNAs. However, for circPLOD2, we only identified few putatively interacting microRNAs and no consistent effect on the predicted miRNA target genes. Based on these observations, we concluded that circPLOD2 is unlikely to interfere with miRNA regulation. Instead, our data indicated that circPLOD2 influences *KLF4* mRNA stability. How this occurs mechanistically requires further studies. One scenario would be a RNA:RNA interaction between *KLF4* mRNA and circPLOD2 that could potentially influence *KLF4* mRNA stability, as recently described for the interaction between circZNF609 and *CKAP5* mRNA^28^. For instance, circPLOD might interfere with RNA-binding proteins and thereby indirectly modulate *KLF4* half-life. Alternatively, circPLOD2 could alter RNA folding, resulting in structural changes in *KLF4* mRNA.

At the transcriptional level, we found that circPLOD2 depletion triggers massive changes in gene expression in pericytes. These changes are mirrored in the hypoxia response when circPLOD2 is strongly upregulated, indicating that the induction of circPLOD2 substantially contributes to the hypoxic response by controlling gene expression. Importantly, using a combination of transcriptional and epigenomic features, we were able to attribute the observed changes to the regulation of the transcription factor KLF4. Using active open-chromatin regions from pericytes that are regulatory elements of circPLOD2-regulated genes, we could predict the DNA binding of all expressed TFs. A direct involvement of KLF4 is evidenced by (i) its downregulation upon circPLOD2 depletion and a corresponding upregulation upon hypoxia when circPLOD2 is induced, (ii) the enrichment and widespread occurrence of KLF4 binding sites in the regulatory elements associated with circPLOD2-regulated genes, and (iii) the accessibility of KLF4-associated regulatory elements, indicating that KLF4 is active under normal conditions in pericytes. Together, these observations suggest that KLF4 targeting via circPLOD2 could be responsible for the massive regulation in gene expression under hypoxia and in response to circPLOD2 depletion. Of note, many of the differentially regulated target genes are involved in cell cycle regulation, suggesting that the increased proliferation of circPLOD2-depleted pericytes could be mediated by the decreased level of KLF4, a well-known cell cycle inhibitor that controls the expression of cell cycle check points (reviewed in ^29^).

KLF4 has already been investigated in different cell types in the vascular system, such as in the perivascular cells, resulting in different and sometimes seemingly contradictory observations. For instance, the described role in the smooth muscle cells seems to be very complex by facilitating phenotypic switching^30^, concomitant that the roles within the smooth muscle cell phenotypes still need to be pinpointed. Similarly, KLF4 in perivascular cells was described to have either beneficial or detrimental effects. On the one hand, *KLF4* knockout in human pericytes and smooth muscle cells induced pericyte bridges, which might be a sign of pericyte detachment^24^. Moreover, smooth muscle cell-specific deletion of *Klf4* in mice reduced the coverage of smooth muscle cells along the resistance arteries and exhibited increased permeability^23^. However, in models of obesity-associated inflammation, smooth muscle cells and pericytes from *Klf4* knockout mice showed a marked protection against vascular inflammation and decreases in proinflammatory macrophages in adipose tissue^31^. Genetic inactivation of *KLF4* in perivascular cells furthermore decreased formation of a pre-metastatic niche and metastasis^32^. Given that the circRNA host gene *PLOD2* is essential for the invasion and metastasis and that circPLOD2 was shown to augment proliferation of ovarian cancer cells^33^, one might speculate that silencing of circPLOD2 could reduce *KLF4* expression to limit cancer progression. Together, these data show that further studies are necessary to elicit the role of *KLF4* in perivascular cells, particularly in pericytes.

Together, our study reports a novel function of circPLOD2 in the control of pericyte functions *in vitro*. The understanding of the noncoding regulatory network is important to understand the regulation of pericyte functions under pathological conditions and may help to identify interventions to improve vessel growth and stability.

## Methods

### Cell culture and growth conditions

Human brain vascular pericytes (passages 2-6; #1200, ScienCell) were cultured in Dulbecco’s Modified Eagle Medium Glutamax (#31966-021; Thermo Fisher Scientific), supplemented with 10% fetal bovine serum (FBS) (#CC-4101; Lonza) and 100 units/ml penicillin and 100 μg/ml streptomycin (#11074440001; Roche). Human umbilical vein endothelial cells (HUVECs; passage 2-3) were purchased from PromoCell (#C-12203) and cultured in EBM Medium from Lonza (#CC-3121), supplemented with hydrocortisone (#CC-4035C; Lonza), ascorbic acid (#CC-4116C, Lonza), bovine brain extract (#CC-4092C; Lonza), epidermal growth factor (#CC-4017; Lonza), 10% fetal bovine serum (FBS) (#CC-4101; Lonza) and Gentamicin/Amphotericin-B-1000 (#CC-4081C; Lonza). All cells were kept at 37 °C and 5% CO_2_. For the pericytes, all cell culture dishes and flasks were coated with 1% Gelatin (#G1393; Merck, in H_2_O) for 30 min at 37 °C. Hypoxia (1% O_2_, 5% CO_2_, 24 h) was induced by using a hypoxic incubator (Labotect, Germany). Cell culture medium was pre-equilibrated overnight. Control cells were incubated in the normal incubator at 21% O_2_, 5% CO_2_ at 37 °C. The hypoxic pO_2_ levels were verified by measuring *VEGFA* mRNA expression and pO_2_ levels with a hypoxia sensing probe from Oxford Optronix (Oxford, UK). For defined cell numbers, cells were washed with Dulbecco’s phosphate-buffered saline (DPBS; #14190-094; Thermo Fisher Scientific) and pericytes detached with 2.5% trypsin (#15090-046; Thermo Fisher Scientific) and endothelial cells with 0.05% trypsin (#25300062; Thermo Fisher Scientific) for 3 min at 37 °C. The enzymatic reaction was stopped with medium and cells were subsequently counted by using a Neubauer Chamber (cells*10^4^/ml). To assess the cell number in Figure 5D, cells from a 6 cm dish with a growth area of 21 cm^2^ was counted.

### RNA extraction and sequencing

RNA deep sequencing was performed by analyzing ribosomal-depleted total RNA of siRNA treated human pericytes (control siRNA or siRNA targeting circPLOD2). Cells were lysed with Qiazol (#79306; Qiagen) for 5 min. The RNA was isolated using the RNeasy Mini Kit (#217004; Qiagen) according to the manufacturer’s instructions, with an additional DNase-I (#79254; Qiagen) digestion for 15min at room temperature. Sequencing was performed on the NextSeq2000 instrument (Illumina) using v3 chemistry, 1×72bp single end setup.

### Preprocessing of RNA-seq data

The overall quality of the RNA-seq data was checked using FastQC (version 0.11.9)^34^. For the trimming of the sequencing reads, flexbar (version 3.5.0)^35^ was utilized with default settings and –min-read-length 20.

### circRNA detection and quantification

circRNAs were detected and quantified with the workflow Calcifer (https://github.com/ZarnackGroup/Calcifer) which can detect and analyze circRNAs from single or multiple datasets. As a starting point, Calcifer uses the output of CIRCexplorer2 (version 2.3.8)^36^ and CIRI2 (version 2.0.6)^37^. For CIRCexplorer2, RNA-seq reads were mapped against the reference genome (GENCODE release 38, GRCh38.p13) using STAR (version 2.7.6a)^38^ with the following parameters: --outFilterMultimapNmax 1 --outFilterMismatchNmax 2 -- alignSJDBoverhangMin 15 --alignSJoverhangMin 15 --chimSegmentMin 15 --chimScoreMin 15 --chimScoreSeparation 10 --chimJunctionOverhangMin 15. The chimeric junctions from STAR were used as input for CIRCexplorer2. For CIRI2, RNA-seq reads were aligned against the same reference genome using bwa (version 0.7.17)^38^ with the parameter -T 19 to pre-filter the resulting alignments. CIRCexplorer2 and CIRI2 were run with default parameters. The circRNAs detected by these tools were then combined and filtered: (i) circRNAs that lacked a canonical splice site or spanned more than 100 kb were filtered out, as described before^11^. To harmonize the results between both tools and achieve comparable quantifications, we recounted the supporting reads for each circRNA from the chimeric junctions reported by STAR. Only circRNAs that had at least two unique reads (i.e., distinct CIGAR strings) across the back-splice junction in at least one replicate were kept as trustworthy. The same mapping of two reads in two different replicates counted as separate uniquely mapped reads. The output of Calcifer specifies the number and position of circRNAs for the individual replicates as well as for the entire dataset. The putative circRNA sequence was reconstructed from the information of the back-splice junction as well as all encompassed annotated exons. To estimate the relative abundance of circRNAs over the corresponding linear transcripts, the circular-to-linear ratio was determined for each circRNA. This was achieved by dividing the number of back-splice reads supporting a given circRNA by the number of reads including linear junctions from the same splice sites, taking the mean count of linear junctions from both splice sites involved in circRNA formation^11^. For the analysis of alternative circPLOD2 isoforms, reads including the back-splice junction of circPLOD2 were taken from the Chimeric.out.junction output of STAR. The CIGAR strings of these reads were used to decoded the possible inclusion of further exons in addition to exons 2 and 3, revealing the occasional inclusion of two intervening, noncoding exons.

### Differential gene expression analysis for circRNAs and linear genes

For the analysis of linear gene expression, aligned reads were counted into the exons of all annotated genes (GENCODE release 38, GRCh38.p13) using htseq-count (version 0.11.3)^39^. These were combined with the raw back-splice read counts for circRNAs that were calculated by Calcifer. Differential gene expression was analyzed using DESeq2 (version 1.34.0)^39^ without independent filtering. Only genes and circRNAs with a mean normalized read count > 1 were considered. Heatmaps were created with ComplexHeatmap (version 2.10.0)^39^ and show z-scores that were computed based on the normalized read counts in all replicates. Coverage tracks for linearly mapped reads, normalized to counts per million (CPM), were generated using bamCoverage from deepTools^40^.

### Gene ontology (GO) enrichment analysis

GO Biological Process (C5 BP) enrichment for differentially expressed genes upon hypoxia and circPLOD2 depletion was tested using the R package hypeR (version 1.10.0)^41^ with an adjusted *P* value < 0.1. The enrichment was tested on all C5 BP annotations provided by the R package msigdbr (version 7.5.1)^42^.

### Identification of significantly enriched transcription factor (TF) binding sites in regulatory elements (REMs)

To identify TFs that could mediate the gene expression changes upon circPLOD2 depletion, we performed a motif enrichment analysis on regulatory elements (REMs) linked to the observed differential expressed genes (DEGs). Using the EpiRegio database (version v1)^43^, a comprehensive resource of more than 2.4 million REMs associated to human target genes, we extracted 68,087 REMs for the top 1000 DEGs after circPLOD2 depletion (sorted by adjusted *P* value, Benjamini-Hochberg correction, from DESeq2). We sorted the REMs in decreasing order based on their DNase I signal in human brain pericytes and excluded those with a mean DNase I signal less than 0.1 (computed with STITCHIT’s REMSelect function^21^), yielding a total of active 16,901 REMs for further analysis. The DNase-seq data was taken from the ENCODE portal^44,45^ with the identifier ENCFF353QSZ. For the 16,901 REMs, the DNA sequence was obtained using bedtools’ getfasta functionality (version 2.27.1)^46^ for the human genome version hg38. 633 human TF binding motifs from the JASPAR database (version 2020)^47^ were downloaded and filtered for those, for which the corresponding TF gene was expressed in our RNA-seq data (transcript per million [TPM] > 1), retaining in 292 TF motifs. The motif enrichment analysis was done using the rank-based PASTAA algorithm^22^ which results in 95 significant TFs (adjusted *P* value ≤ 0.01, Benjamini-Hochberg correction). Using FIMO (version meme-5.2.0)^48^, the binding sites of the significant TFs were determined for each REM. Next, we counted the predicted TF hits per REM and visualized the counts as heatmap using the R package complexHeatmap (R version 2.10.0). For the analyses described, we slightly modified the following workflow from our epiregio github repository:https://github.com/TeamRegio/ApplicationScenarioExamples/tree/master/PASTAA_Fimo_analysis.

### Computation of DNase I sensitivity signals

To compute the DNase I signal of REMs with at least one predicted KLF4 binding site (3,025 of the 68,086 REMs), we used deepTools^40^. We extracted the REM positions and used them together with the DNase I signal as input for the following two commands: computeMatrix reference-point -S <DNase-signal.bigWig> -R <REMs.bed> -a 2000 -b 2000 -o <output.tab.gz> --referencePoint center; plotHeatmap -m <output.tab.gz> -o <heatmap.pdf> --regionsLabel REMs.with.KLF4.binding.site --legendLocation none --samplesLabel DNase-signal -x ‘distance (bp)’.

### miRNA target site predictions

miRNA target sites in circRNAs were predicted as part of the Calcifer workflow. Initially, to detect potential miRNA target sites, the exonic sequence was determined for each circRNA based on gene annotation (GENCODE release 38, GRCh38.p13), considering all exons between the back-splice junction. To include potential sites across the back-splice junction, the sequence was duplicated and concatenated. All human miRNA input sequences were downloaded as .*fasta* format from miRBase release 22.1 (as accessed on 17.05.2021)^49^. In order to obtain miRNA circPLOD2 interactions, miRanda (version 3.3a) was used with parameters alignment score 80 (-sc 80) and energy threshold -15 (-en 15). For each circRNA, all hits for each miRNA were summed up and divided by two to account for the duplicate sequence. The predicted miRNA binding sites in the circPLOD2 transcript were overlapped with AGO1-4 CLIP binding sites (downloaded from doRiNA 2.0^18^), overlapped using bedtools v2.30.0 intersect^50^.

### qRT-PCR experiments

After RNA isolation (described above), the concentration and quality were determined using the NanoDrop2000 spectrophotometer from Thermo Fisher Scientific. 1 μg of RNA from each sample was reverse-transcribed by M-MLV Reverse Transcriptase using random hexamer primers (master mix; all reagents were purchased from Thermo Fischer Scientific: (I) 0.4 µl dNTPs (#362271), 1 µl random hexamers (#S0142), 1 μg of RNA, filled up to 26 µl with H_2_O and; (II) 8 µl 5x first strand buffer (#18057-018), 4 µl DTT (100mM; (#18057-018), 1 µl RNase Inhibitor (#N808-0119), 1 µl MuLV (#28025-013). The PCR was run with the following program: Mastermix (I): 65 °C 5 min; Mastermix (I+II): 25 °C 10 min, 37 °C 50 min, 70 °C 15 min, 4 °C). The reversed-transcribed RNA was used for real-time quantitative PCR (qPCR) by using Fast SYBR Green Mastermix (#4385612, Life Technologies) on a ViiA7-Real-time qPCR System from Life Technologies with the following master mix: 5 µl Fast SYBR Green Mastermix, 1.5 µl H_2_0, 1 µl forward and reverse human primer mix (10 µM each; *RPLP0*-fw: GGCGACCTGGAAGTCCAACT, *RPLP0*-rev: CCATCAGCACCACAGCCTTM; *circPLOD2*-fw: TGGAACACTATGCTGATCAAGA, *circPLOD2*-rev: GACTGCATAAATCGATGGAATCC; *linPLOD2*-fw: GCAGATGGAATTTTGTGGCC, *linPLOD2*-rev: GGAGCATAGCCAATAAATCCTCC; *VEGFA*-fw: AATGTGAATGCAGACCAAAG, *VEGFA*-rev: GACTTATACCGGGATTTCTTG; *cHIPK3*-fw: TCGGCCAGTCATGTATCAAAGA, *cHIPK3* rev: GACCAAGACTTGTGAGGCCA; *HypERlnc*-fw: AGGCCAGAGGATGGAAAAGG *HypERlnc*-rev: TTTGCATCTCCCAACCAGCA; *KLF4*-fw: TCCCATCTTTCTCCACGTTC, *KLF4-rev*: AGTCGCTTCATGTGGGAGAG; all primers were purchased from Sigma Aldrich). Ct values were normalized against ribosomal *RPLP0* and relative gene expression was determined by the 2^-ΔCt^ method.

### Sanger sequencing

For Sanger sequencing, isolated RNA from pericytes were reverse transcribed with random hexamers. Afterwards, cDNA was amplified by using Expand Long Template Enzyme mix (#11681834001; Roche) with the following master mix: 1.75 µl dNTPs 10 mM, 5 µl 10x PCR buffer with MgCl_2_, 0,75 µl Expand Long Template Enzyme mix, 36 µl H_2_0, 5 µl template, and 1.5 µl primer mix (10 µM each; forward: ATTTATCATTCAGTAAACATGAC; reverse: TATTAGTCATAACTGTAGCAAC) and the PCR program: 94 °C 2 min, 34 cycles of [94 °C 20 s, 55 °C 1 min, 65° C 3:15 min], 65 °C 7 min, 4 °C). The amplified product was cleaned by using the PCR Clean Up Kit (#740609.50; Macherey-Nagel), following the manufacturer’s instructions. 41.58 ng DNA and the same primer were used for sequencing with Microsynth.

### RNase R digestion

For RNase R digestion, 5 µg RNA was incubated with 5 U RNase R (#RNR07250; Epicentre) in 1× RNase R buffer (#RNR07250; Epicentre) in a 10 µl reaction volume at 37 °C for 10 min followed by heat inactivation at 95 °C for 3 min. 1 µg of the digested RNA was reverse-transcribed as described above and validated by PCR using the KOD Xtreme HotStart Polymerase Kit (#71975-3; Merck, PCR program: 94 °C 2 min, 26-32 cycles of [98 °C 10 s, 60 °C 20 s, 68 °C 15 s], 4 °C) by using the following primer: *PDGFR-β*-fw: ACAATGACTCCCGTGGACTG, *PDGFR-β*-rev: CTCGGCATCATTAGGGAGGA; *VEGF-A-*fw: AATGTGAATGCAGACCAAAG, *VEGFA-*rev: GACTTATACCGGGATTTCTTG; *circPLOD2*-fw: GACTGCATAAATCGATGGAATCC *circPLOD2*-rev: TGGAACACTATGCTGATCAAGA) according to the manufacturer’s instructions. Samples were stained with Orange DNA loading dye (#R0631; Thermo Fisher Scientific) and analyzed on an 2% agarose gel (#A9539; Sigma Aldrich), containing MIDORI green (#617004 Biozym); at 160 V for 35 min. O′GeneRuler™ 100bp plus (#SM1153; Thermo Fisher Scientific) was used as ladder.

### Poly(A) RNA enrichment

To fractionate the RNA into RNA with and without poly(A) tail, oligo-d(T)25-Magnetic Beads (S14195, New England Biolabs) were used. Thereby, 5 µg RNA was incubated with washed beads (washing buffer: 300 mM NaCl [#AM9760G; Thermo Fisher Scientific], 20 mM Tris-HCl [#T2694; Sigma Aldrich], 0.01% NP-40 [#A2239; AppliChem]) in binding buffer (300 mM NaCl, 40 mM Tris-HCl, 0.02% NP-40) for 10 min on a shaker (180 mot/min). For separation, samples were placed on a magnet. The supernatant contains the oligo-d(T)_25_-negative RNA. After several washing steps, oligo-d(T)_25_-positive RNA was eluted by incubation in elution buffer (150 mM NaCl, 20 mM Tris-HCl, 0.01% NP-40) for 2 min at 50 °C.

### siRNA-mediated depletion

For small interfering RNA (siRNA)-mediated RNA depletion, pericytes were seeded in a cell culture dish and incubated till they achieved an 80% confluency. Cells were transfected with control siRNA (siScr, CGUACGCGGAAUACUUCGATT; final concentration: 60 nm), siRNA targeting circPLOD2 (sicPLOD2. UACUGAAUGAUAAAUUAUUTT, final concentration of 60nM) linear PLOD2 (linPLOD2 GGCUUCAUUUUAUCCGGGATT; final concentration: 40 nM) by using Lipofectamine RNAiMax (#L13778; Thermo Fischer Scientific) in Opti-MEM-I GlutaMAX (#51985-026; Life technologies), according to the manufacturer’s instructions. All siRNAs were purchased from Sigma Aldrich. After 4 h, the transfection mix was replaced by cell culture medium.

### Lentiviral GFP expression

Lentivirus stocks were produced in HEK293T cells, cultured in DMEM GlutaMAX (#10566-016; Invitrogen), containing 10% FBS and 100 units/ml penicillin and 100 μg/ml streptomycin (#11074440001; Roche) with the packing plasmids pCMV∆R8.91 and pMD2.G and a plasmid SEW carrying the GFP-ORF as previously described^51^. Pericytes were transduced with the lentivirus for 24 h and GFP expression was confirmed by fluorescence microscopy.

### *KLF4* overexpression

To overexpress *KLF4* and simultaneously perform a knockdown of circPLOD2, Lipofectamine 3000 (#L3000; Thermo Fischer Scientific) in Opti-MEM-I GlutaMAX (#51985-026; Life technologies) was used, according to the manufacturer’s instructions. Cells were transfected either with a mock plasmid and a control siRNA (= control; final concentration of the plasmid: 0.4 µg; siRNA: 80 nM) or a plasmid containing *KLF4* sequence and siRNA targeting circPLOD2 (KLF4 OE+sicircPlod2, final concentration of the plasmid: 0.4 µg; siRNA: 80 nM). After 4 h, the transfection mix was replaced by cell culture medium. The following plasmids were used: pCMV6-XL5-KLF4 (#SC123501; Origene) and pCMV6-Entry (#PS100001; Origene).

### Tube formation assay

For the tube formation assay, HUVECs stimulated with pericyte supernatant were used. Pericytes were transfected with siRNAs (siScr or sicPlod2) as described above for 24 h. The supernatant was collected, centrifuged at 700 x g for 5 min and mixed with HUVEC medium. HUVECs were treated for 24 h. For the assay, a 48-well plate was coated with 150 µl Matrigel per well (growth factor-reduced, #354230, BD Pharmingen) for 30 min at 37 °C before adding 100,000 HUVECs in 250 µl medium with the supernatant of either siScr-treated or sicPlod2-treated pericytes. The cells were incubated for 24 h at 37 °C, 5% CO_2_. Five randomly chosen fields per view per Matrigel assay were acquired using Axiovert1000 microscope (Zeiss; Axiovision 4.8 Software) and the tube length and number of nodal were measured by using Zen3.4 lite Software.

### Migration assay

To analyze the migration, the Boyden chamber assay was used. 50,000 transfected pericytes (siScr or sicPlod2) were seeded into a 10% gelatin-coated 8 µm membrane Boyden chamber (#351152; Corning), placed in a 24-well plate (#353504; Corning). After 3 h of incubation, the cells were fixed with 4% paraformaldehyde (#28906; Thermo Fisher Scientific) for 10 min at room temperature. Cells were stained with DAPI (1:1000; #4083; Cell Signaling) in DPBS for 10 min. The bottom of the Boyden chamber was analyzed. Five randomly chosen fields per view per condition were acquired using a TCS-SP8 microscope (Leica) and the number of migrated pericytes was determined by counting DAPI-positive nuclei with ImageJ.

### Matrigel co-culture assay

GFP-expressing pericytes and Dil-LDL stained HUVECs were used for the co-culture assay. Pericytes were transfected with siRNAs and/or plasmids, as described above, for 48 h. HUVECs were stained with acetylated Dil-LDL overnight (10 µg/ml; #L3484, Invitrogen) for 24 h. For the assay, 48-well plate was coated with 150 µl Matrigel per well (growth factor-reduced, #354230, BD) for 30 min at 37 °C before adding 100,000 HUVECs in 500µl EC-medium. After 3 h, 10,000 pericytes in 500 µl PC-medium were added and incubated for additional 3 h. The medium was removed and a second layer of 150 µl Matrigel was added carefully on top. After 30 min, 500 µl cell medium (equal amounts of Pericyte and HUVEC medium) was added. After 16 h, the assay was fixed for 10 min with 4% paraformaldehyde (#28906; Thermo Fisher Scientific), and 3-8 randomly chosen fields per view per Matrigel assay were acquired using confocal Z-stack imaging (TCS-SP8; Leica). Attached pericytes were counted by using the LASX Software.

### Bromodeoxyuridin (BrdU) incorporation assay

Pericytes were cultured on a gelatin coated glass 8-well chamber slide (#177402, Thermo Fisher Scientific). After 24 h the cells were treated with control siRNAs (siScr) or siRNAs targeting circPLOD2 (sicPlod2) for additional 24 h. Afterwards, cells were treated with 10 nM BrdU or DPBS as control for 6 h at 37 °C. The cells were fixed with ice-cold methanol for 15 min at 4 °C, followed by three washing steps with DBPS. The DNA was hydrolyzed by using 2 M HCl (#J/4315/15, Fisher Chemicals) for 45 min at 37 °C, washed four times with Tris buffered saline (#sc24951, ChemCruz) on shaker. Cells were blocked with 10% donkey serum (#ab7475; Abcam) in DPBS for 30 min at room temperature and stained with anti-BrdU-488 (1:50, #51-23614L, BD Pharming) and HOECHST (1:500; #5117, Tocris) in blocking buffer at 4 °C, overnight. On the next day, cells were washed three times with DPBS containing 0.01% Tween-20 (#A139, Applichem) and slides were mounted (#71-0016, KPL). Three randomly chosen fields per view per condition were acquired using confocal imaging (TCS-SP8, Leica) and positive BrdU cells were counted, normalized to all cells in the field of view.

### Cell viability assay

To measure the cellular viability, the RealTime-Glo™ MT Cell Viability assay (#G9711; Promega) was used, according to the manufacturer’s instructions. Briefly, pericytes were transfected as described above and after 24 h, 4,000 cells were seeded into a 96-well plate. Cells were treated with the NanoLuc enzyme and substrate, and the plates were sealed with a breathseal foil (#676050; Greiner). The luminiscence (RLU) was measured after 48 h, by using the GloMax plate reader (Promega).

### Actinomycin D assay

Transfected pericytes were treated with 1 µg/ml Actinomycin-D (#A9415; Sigma Aldrich) or DMSO (1:1000, #A994.2; Roth) for 1 h. Afterwards cells were lysed in Qiazol and RNA was isolated as described above. mRNA levels were measured by qRT-PCR and analyzed with the 2^-Ct^ method.

### Statistics

Gaussian distribution was calculated by using Shapiro Wilk normality test. Mann–Whitney U-test or Student’s *t*-test were used to test for statistical differences between two groups as appropriate. For more group’s analysis of variance (ANOVA) with multiple comparison test and Tukey or Bonferroni correction was used. Outlier were identified by using Grubbs outlier test. A *P* value < 0.05 was considered as statistically significant. GraphPad Prism 9 were used to calculate statistical differences.

## Data availability

Published data for human brain vascular pericytes in normoxic and hypoxic condition were taken from Gene Expression Omnibus (GEO; www.ncbi.nlm.nih.gov/geo/) accession number GSE84730^16^. DNase I sensitivity data for human pericytes were downloaded from the ENCODE portal (https://www.encodeproject.org/) with the identifiers ENCFF353QSZ. The RNA-seq data generated in this study will be deposited in the NCBI Gene Expression Omnibus.

## Acknowledgement

The authors would like to thank Astrid Wiederer and Florian C Bischoff for technical support and all members of the Zarnack lab for discussion. We acknowledge the ENCODE Consortium and the laboratory of John Stamatoyannopoulos (University of Washington) for providing the DNase-seq data for human pericytes.

## Funding

The study was supported by the German Research Foundation (DFG), TRR 267 (Project ID 403584255) Project A01 to S.D. and K.Z, and Project A04 to M.S.L., and Project Z03 to M.H.S..

## Supplementary Figures

**Figure S1:**
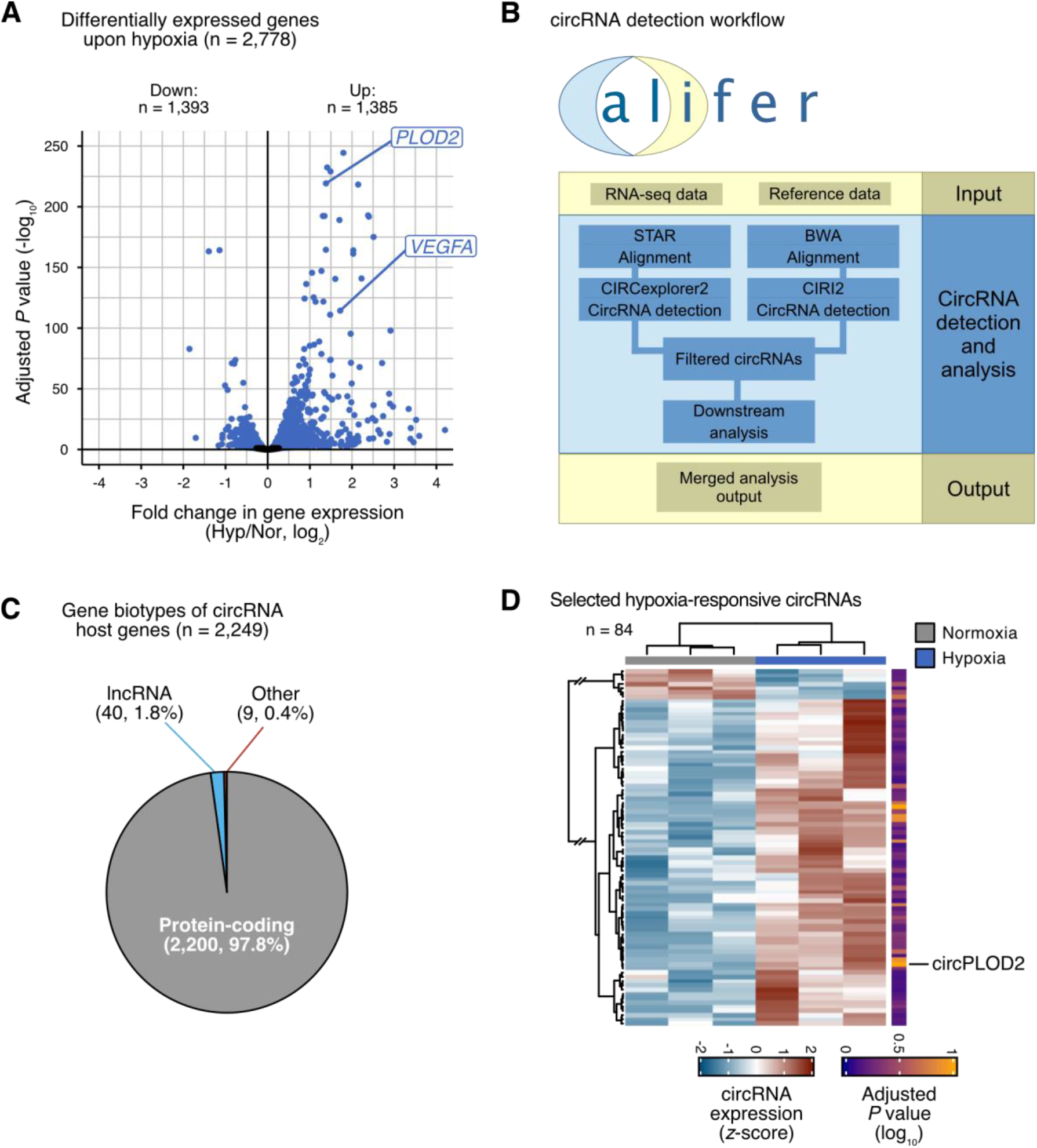
Linear transcripts and circRNAs change upon hypoxia. **(A)** 2,778 genes significantly change in expression after hypoxia. Volcano plot shows log_2_-transformed fold change in gene expression (Hyp/Nor) against adjusted *P* value. Significantly regulated genes are shown in blue (adjusted *P* value<0.1, Benjamini-Hochberg correction). **(B)** Overview of Calcifer workflow to detect, quantify and analyze circRNAs. circRNA detection is based on the output from CIRCexplorer2 and CIRI2, which is further processed and filtered. The identified circRNAs are then subjected to downstream analyses, including microRNA (miRNA) target site prediction. **(C)** Most circRNAs originate from protein-coding genes. Pie chart shows gene biotype of 2,249 genes that harbor 6,187 detected circRNA in human brain vascular pericytes. of all circRNAs. Total numbers and percentages are given. **(D)** Most hypoxia-responsive circRNAs are upregulated. Heatmap shows normalized read counts (*z*-scores) for 84 hypoxia-responsive circRNAs (coefficient of variation < 1, adjusted *P* value < 0.8, DESeq2). Color scale (right) indicates negative log_10_-transformed adjusted *P* value. circPLOD2 is indicated on the right.

**Figure S2:**
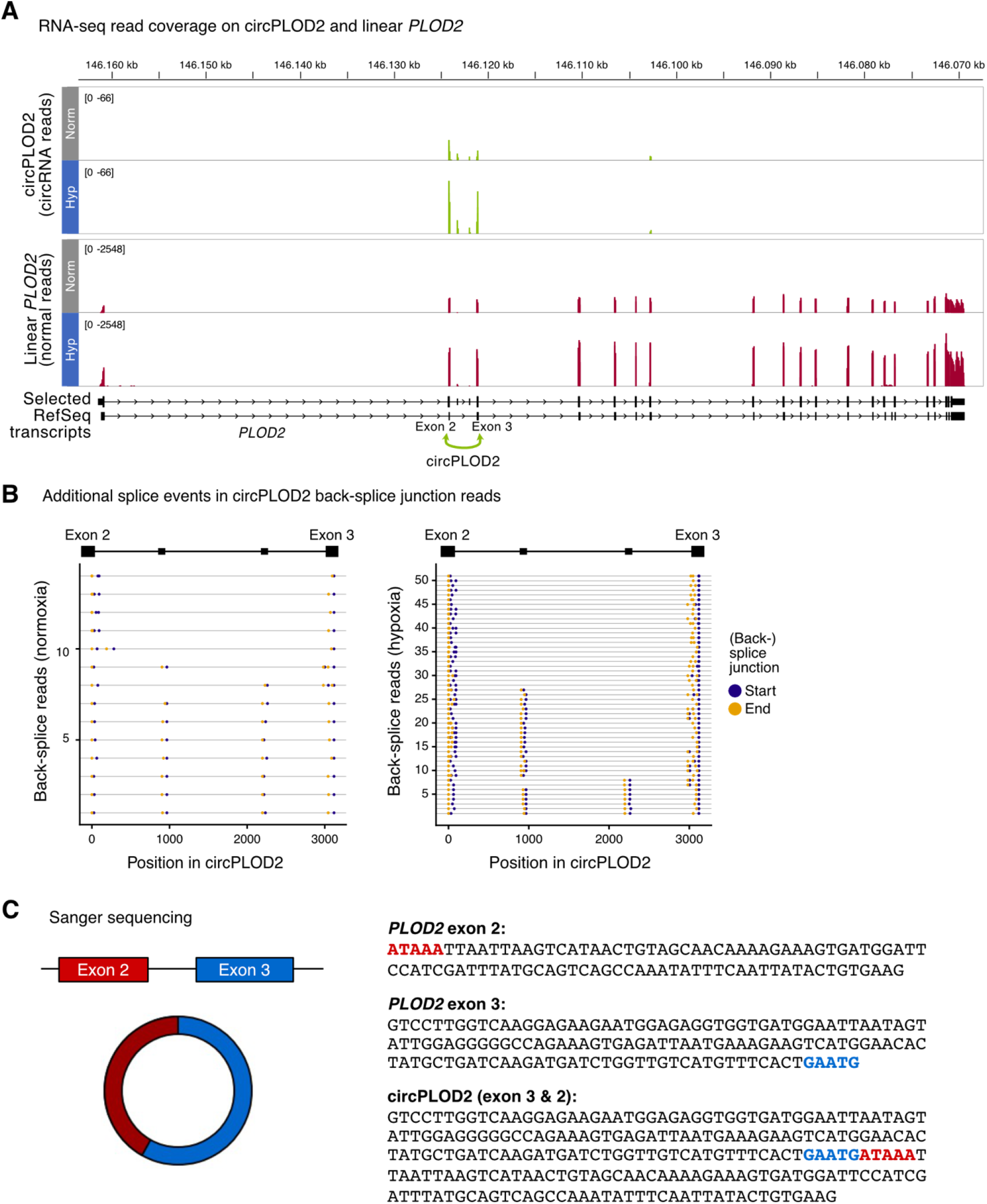
circPLOD2 is generated via back-splicing between *PLOD2* exon 2 and 3. **(A)** Genome browser view depicts coverage of circRNA reads (back-splice junctions, green) and linear reads (darkred) for *PLOD2* gene. The circPLOD2 isoform harboring only exons 2 and 3 is the major isoform from pericytes in normoxia (Nor, grey) and hypoxia (Hyp, blue). Further circPLOD2 isoforms additionally include two intervening, noncoding exons. Within each read type, coverage tracks for normoxia and hypoxia are shown on the same scale. **(B)** Back-splice reads inform about internal alternative splicing events within circPLOD2. Plot depicts detected splice junctions in all back-splice read pairs (rows) for circPLOD2 in normoxia (left) and hypoxia (right). A fraction of reads shows additional inclusion of two intervening exons between exons 2 and 3. **(C)** The back-splice site and sequence of circPLOD2 was confirmed by Sanger sequencing (n=2). The main circPLOD2 isoform consists of exon 2 (92 nt, red) and exon 3 (137 nt, blue), resulting in a circRNA with a total length of 229 nt (left). The back-splice site is highlighted.

**Figure S3:**
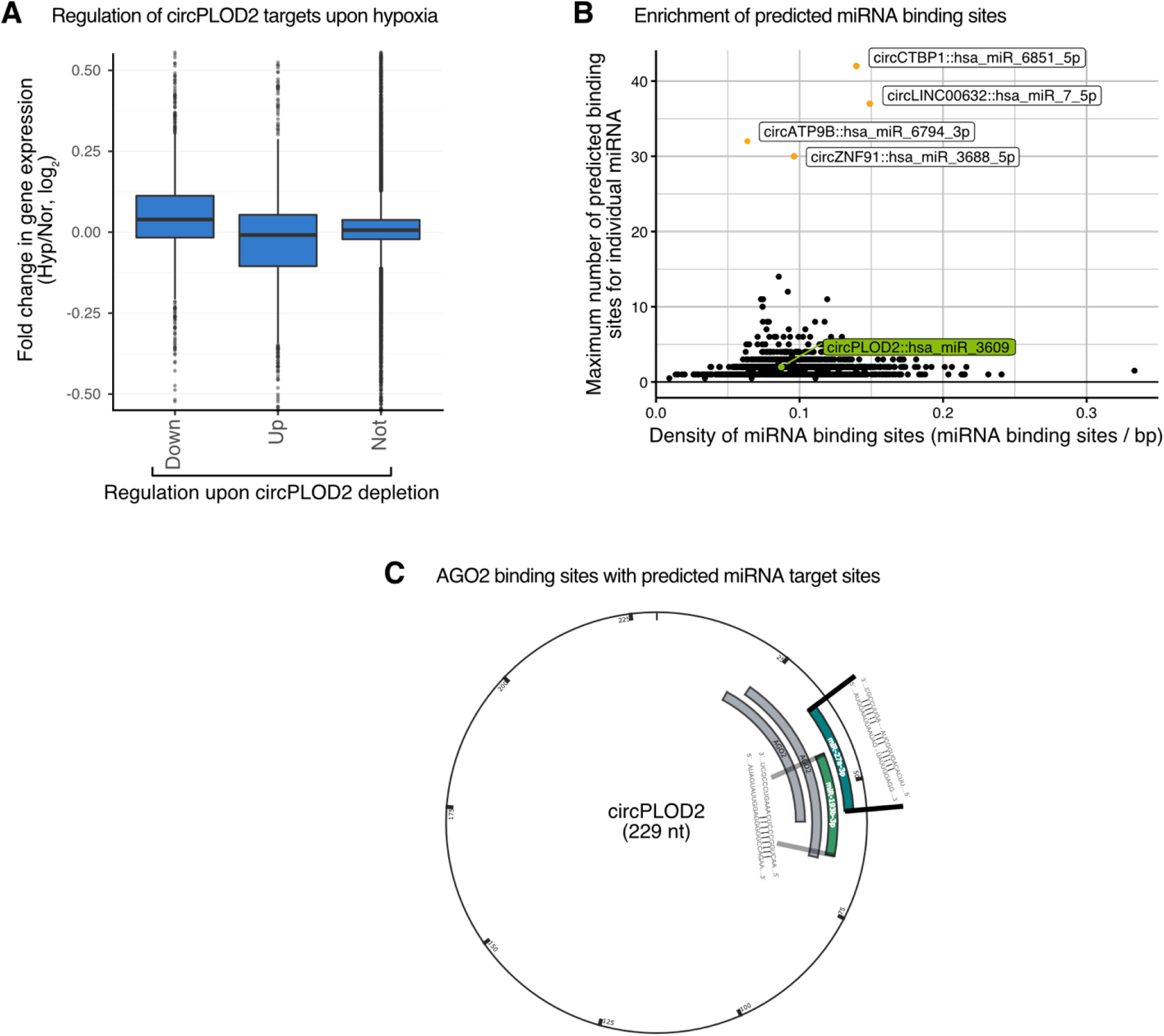
Regulation of circPLOD2 targets upon hypoxia and potential interaction of circPLOD2 with microRNAs (miRNAs). **(A)** circPLOD2-regulated genes respond in hypoxia when circPLOD2 is upregulated. Boxplot shows shrunken log_2_-transformed fold changes upon hypoxia for genes that are downregulated (n = 1,060), upregulated (n = 1450) and not regulated (n = 21,748) upon circPLOD2 depletion. **(B)** circPLOD2 is not enriched for miRNA target sites. Scatter plot shows the density of miRNA target sites against the maximum number of predicted targets sites for any individual miRNA for circRNAs found in pericytes (n = 3,708). circRNAs with > 25 miRNA target sites for at least one individual miRNA are labeled. **(C)** circPLOD2 harbors two AGO2 binding sites with predicted miRNA target sites. By using publicly available AGO2 pulldown data, two AGO2 binding sites overlapping with two predicted miRNA binding sites (miR-27a-3p and miR-193b-3p) on circPLOD2 were identified. The predicted target genes for both miRNAs (targetScan 8.0 https://www.targetscan.org/vert_80/) showed no directed regulation in circPLOD2-depleted pericytes (no significant enrichment for genes regulated in either direction upon circPLOD2 depletion, Fisher’s exact test).

## Notes

### Competing Interest Statement

The authors have declared no competing interest.

